# NanoBERTa-ASP: Predicting Nanobody Binding Epitopes Based on a Pretrained RoBERTa Model

**DOI:** 10.1101/2023.09.29.560264

**Authors:** Shangru Li, Xiangpeng Meng, Rui Li, Bingding Huang, Xin Wang

## Abstract

Nanobodies, also known as VHH or single-domain antibodies, are a unique class of antibodies that consist only of heavy chains and lack light chains. Nanobodies possess the advantages of both small molecule drugs and conventional antibodies, making them a promising class of therapeutic biopharmaceuticals. Studying the characteristics of nanobody sequences can aid the development and design of nanobodies. An important challenge in this field is accurately predicting the binding sites between nanobodies and antigens. The binding site is the region where the nanobody interacts with the antigen. The precise prediction of these binding sites is crucial for comprehending the interaction mechanism between the nanobody and the antigen, facilitating the design of effective nanobodies, as well as gaining valuable insights into the properties of nanobodies.

Nanobodies typically have smaller and more flexible binding sites than traditional antibodies, however predictive models trained on traditional antibodies may not be suitable for nanobodies. Moreover, the limited availability of antibody- derived nanobody datasets for deep learning poses challenges in constructing highly accurate predictive models that can generalize well to unseen data.

To address these challenges, we propose a novel nanobody prediction model, named NanoBERTa-ASP (Antibody Specificity Prediction), which is specifically designed for predicting nanobody-antigen binding sites. The model adopts a training strategy more suitable for nanobodies by leveraging an advanced natural language processing (NLP) model called BERT (Bidirectional Encoder Representations from Transformers). The model also utilizes a masked language modeling approach to learn the contextual information of the nanobody sequence and predict its binding site.

The results obtained from training NanoBERTa-ASP on nanobodies highlight its exceptional performance in Precision and AUC, underscoring its proficiency in capturing sequence information specific to nanobodies and accurately predicting their binding sites. Furthermore, our model can provide insights into the interaction mechanisms between nanobodies and antigens, contributing to a better understanding of nanobodies, as well as accelerating the development and design of nanobodies with potential therapeutic applications. To the best of our knowledge, NanoBERTa-ASP is the first nanobody language model that achieved high accuracy in predicting the binding sites.

**Github repository:** https://github.com/WangLabforComputationalBiology/NanoBERTa-ASP

## 1. Introduction

Antibodies are vital components of the human immune system, characterized by their exceptional specificity and high affinity. They have extensive applications in disease diagnosis, treatment, and prevention. Nanobodies, a unique class of small antibody molecules, differ from conventional antibodies in that they naturally lack light chains.[1] This inherent feature renders nanobodies less prone to mutual adhesion and aggregation. However, their variable heavy chain (VHH) region exhibits structural stability and antigen binding activity comparable to that of full-length antibodies. Nanobodies are considered the smallest functional units known to bind target antigens. Nanobodies possess the advantages of both conventional antibodies and small molecule drugs.[2] Nanobodies are increasingly being recognized as a promising class of therapeutic biopharmaceuticals in the field of therapeutic biomedicine and clinical diagnostic reagents.[3] However, the design and development of nanobodies remain a challenging issue, requiring the resolution of numerous technical hurdles. One key challenge is accurately predicting the binding epitopes between nanobodies and antigens. The binding epitope refers to the region where nanobodies interact with antigens. Accurate prediction of the binding epitopes is crucial for understanding the interaction mechanisms between nanobodies and antigens, as well as for designing improved nanobodies.[4] Studying the characteristics of nanobodies and accurately predicting their binding epitopes holds significant importance.

In recent years, with the advancements in artificial intelligence and deep learning technologies [5], training antibody models using large-scale antibody data has emerged as a novel approach for antibody design and optimization. Compared to traditional antibody research methods, deep learning techniques offer reduced time and cost requirements. With the assistance of computers, antibody researchers can handle larger datasets, predict the properties and functions of unknown antibodies, significantly improving the accuracy of antibody research. Currently, the mainstream methods in deep learning for antibodies are language models and graph neural network models. Graph neural network models can learn the relationships between antibody residues and represent the structure of epitopes, enabling tasks such as antibody docking, pairing, and epitope prediction.[6] Language models, on the other hand, can learn sequence data from a large volume of data, facilitating tasks such as antibody sequence generation, recovery, and epitope prediction.

In this work, we developed a training model called NanoBERTa-ASP, which achieved outstanding results of nanobody on smaller training datasets compared to other models. Our pre-training dataset consisted of approximately 30 million human heavy chain BCR sequences. The finetuning dataset comprised around 2,200 annotated examples, including 1,300 nanobody annotations and 900 antibody heavy chain annotations. NanoBERTa-ASP was built upon the model architecture of RoBERTa, a widely used generalized model. While RoBERTa was initially designed for handling textual data, antibody sequences are also composed of strings of amino acids. Therefore, it is feasible to apply the RoBERTa model to analyze and predict antibody data. In recent studies, researchers have successfully employed RoBERTa in the field of antibody research, achieving promising results. These endeavors demonstrate the potential utility of RoBERTa in antibody-related investigations, showcasing its effectiveness in tasks such as antibody sequence analysis, antigen-antibody interaction prediction, and other relevant studies.

The results demonstrated that NanoBERTa-ASP achieved high accuracy in predicting nanobody binding epitopes, indicating its ability to learn the sequence features of nanobodies. [7]

## II. Experimental procedures

### A. Pretrain dataset

To pretrain NanoBERTa-ASP, we downloaded 70 research-based human unpaired antibody heavy chain sequences from the Observed Antibody Space (OAS) database on April 16, 2023. [8] We removed sequences containing unknown amino acids. To ensure that the model could better capture sequence features, we selected sequences with a minimum of 20 residues in the CDR1 region and a minimum of 10 residues in the CDR3 region. Subsequently, We partitioned the entire collection of 31.01 million unpaired heavy chain sequences into mutually exclusive sets for the purpose of training, validation, and testing. This was done to ensure that there was no overlap between the sequences included in each set.

The pretraining dataset consisted of approximately 24.8 million heavy chain sequences, while the pretrained validation and test sets each contained around 3.1 million heavy chain sequences.

### B. Finetune dataset

we downloaded 7,255 antibody PDB files from The Structural Antibody Database (SAbDab) on April 17th, 2023 to fine-tune our model for binding site prediction.[9] We initially filtered 5,134 crystal structures with an accuracy of 3.0 Å or higher. Structures that were defined as half-antigen binding in IMGT were removed as they did not meet the binding requirements of our nanobodies. Using a distance criterion of less than 4.5 Å as the labeled binding site, we selected 1,070 sequences of labeled nanobodies and 4,400 heavy chain sequences. We then randomly split the data into training, validation, and test sets in an 8:1:1 ratio. To ensure that the model is optimized for the nanobody task, we only used nanobody data in the validation and test sets, while heavy chains were only used to augment the training set. Therefore, we randomly selected a subset of heavy chains to include in the training set.

### C. NanoBERTa-ASP pre-training

NanoBETRa is a pre-trained model based on a modified version of the Roberta model. The vocabulary used for training consists of 24 tokens, including 20 amino acids and 4 identification tokens (<s>, </s>, <mask>, <pad>). The entire sequence is treated as a sentence, with the sequence being identified by the start token <s> and the end token </s>. The MLM(Masked Language Modeling) method was chosen for training, with 15% of the amino acids being perturbed. Similar to the Roberta setup, 80% of the tokens were replaced with <mask>, 10% were replaced with randomly selected amino acids, and 10% were left unchanged. During pretraining, the model predicts the residual of masked positions. for seq in each batch, the loss is:

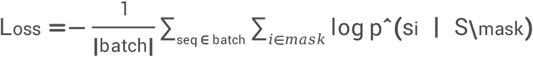

p^^^(si ∣S**\**mask) represents the prediction probability of the model for the label (si) at the i-th position, under the condition that the other parts of the sequence (seq)are removed except for the masked position Mask.

### D. NanoBERTa-ASP fine-tuning

We consider the task of epitope prediction as a binary token classification task, where NanoBERTa-ASP predicts whether each residue in a nanobody sequence is an epitope or not. To achieve this, we add a binary classification head on top of the pre-trained model to label the sequences. During training, the model uses cross-entropy loss function to calculate the difference between the predicted probability p and the true label y, then updates the model parameters using backpropagation algorithm. The loss function used during fine-tuning is:

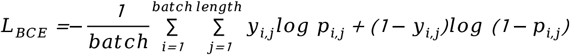

## III. Result

### A. Attention mechanism can focus on the structure of the sequence

As a RoBERTa-based model, NanoBERTa-ASP also has the same multi-head attention mechanism as RoBERTa. The attention heads of NanoBERTa-ASP can focus on different parts of the sequence. NanoBERTa-ASP exhibits a higher degree of attention towards the highly variable CDR3 region. For example, when we input the nanobody Nb-ER19 into the model and output the attention layers in the form of a heatmap, we can observe that the sixth head of the twelfth layer of the model has a special attention on the positions of ASN32 and VAL33 in CDR1, LEU98 in CDR3 (PDB:5f7y) (Fig 2A). After validation using PyMOL, it was found that there was a connection at this position (Fig 2B), and LEU98 is also part of the epitope. This indicates that the model can learn certain structural features of antibodies through the annotated sequences. [10]

**Fig 1.**
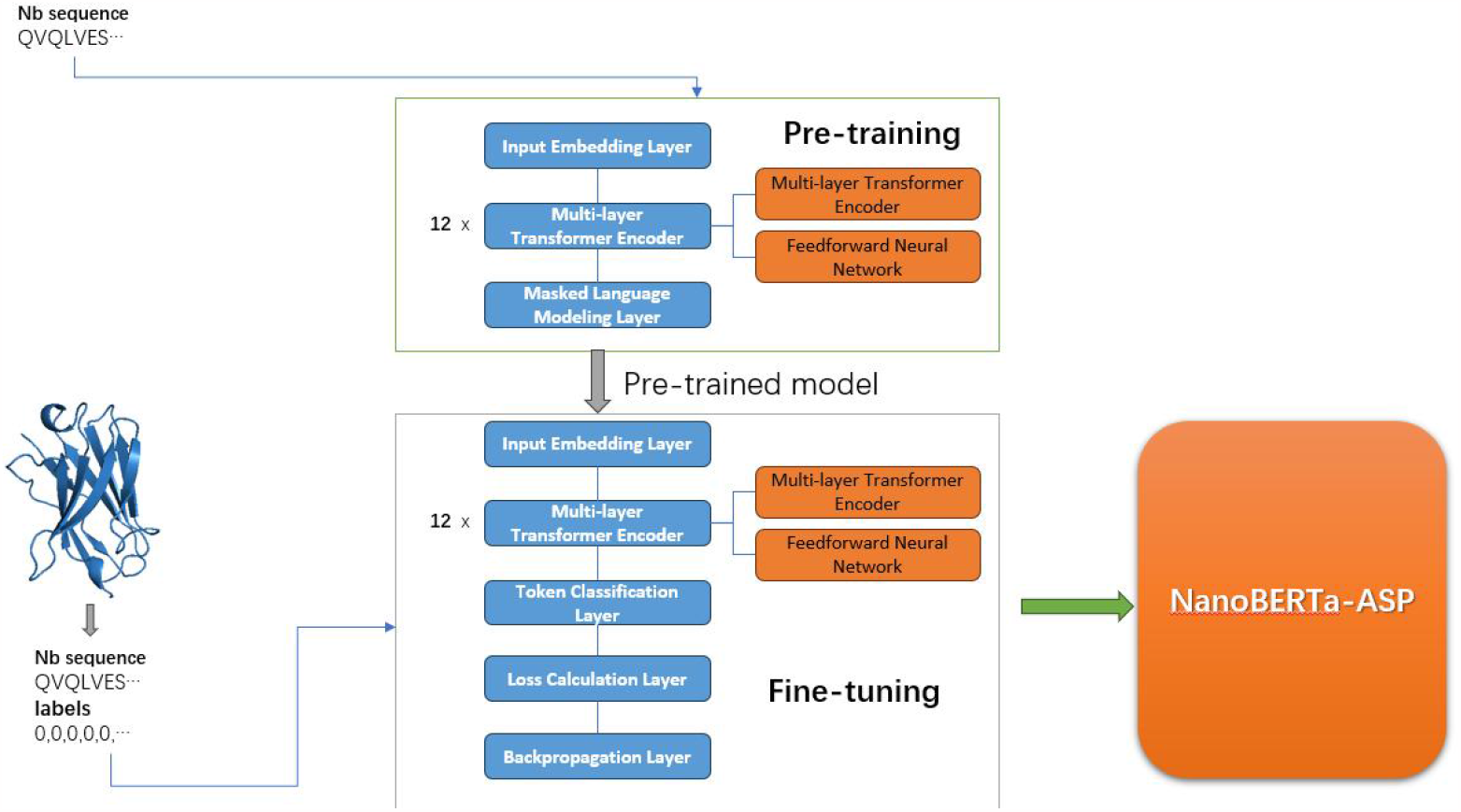
The flowchart of NanoBERTa-ASP.

**Fig 2.**
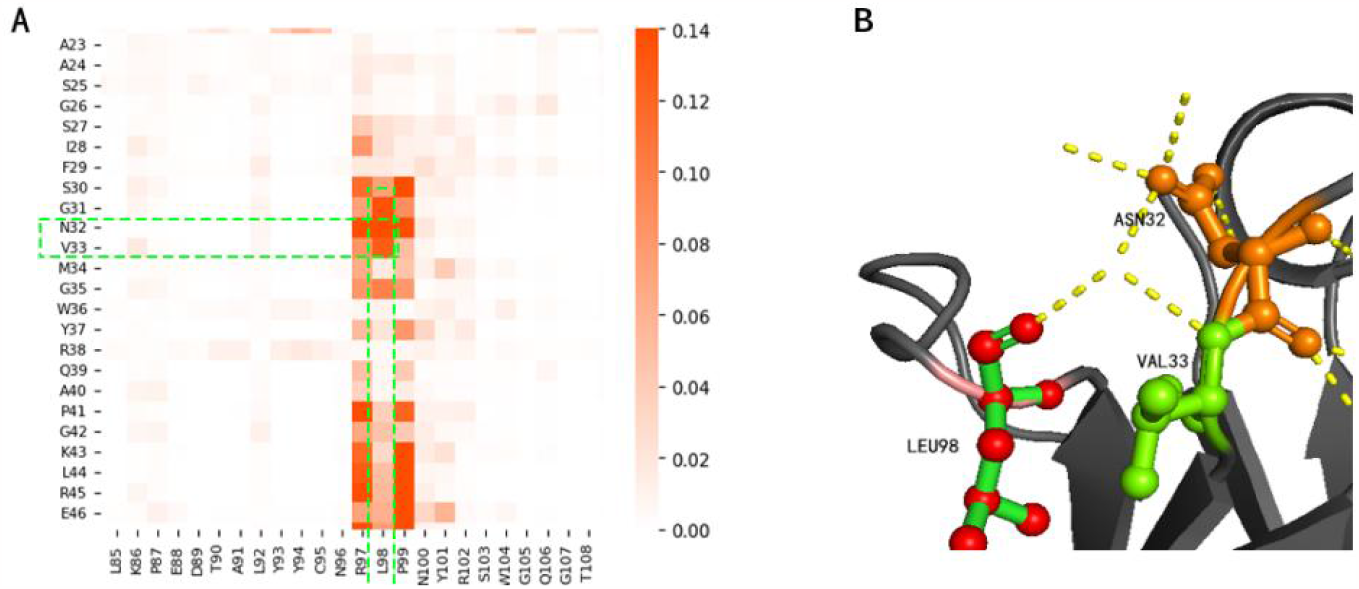
(A) Self-attention heatmap from NanoBERTa-ASP’s 12th layer, sixth head for PDB:5f7y; (B) The schematic diagram of the 3D structure of PDB:5f7y show contacts.

### B. Performance of the NanoBERTa-ASP

Our model can predict binding sites in both CDR and non-CDR regions. Fig 3B and 3D show the predicted binding sites of NanoBERTa-ASP, compared with the computed binding sites from PDB shown in Fig3A and 3C. NanoBERTa-ASP can accurately identify the binding sites on the nanobody sequence, as demonstrated by the comparison with the computed binding sites from PDB (5f7y and 2wzp). [10][11]

**Fig 3.**
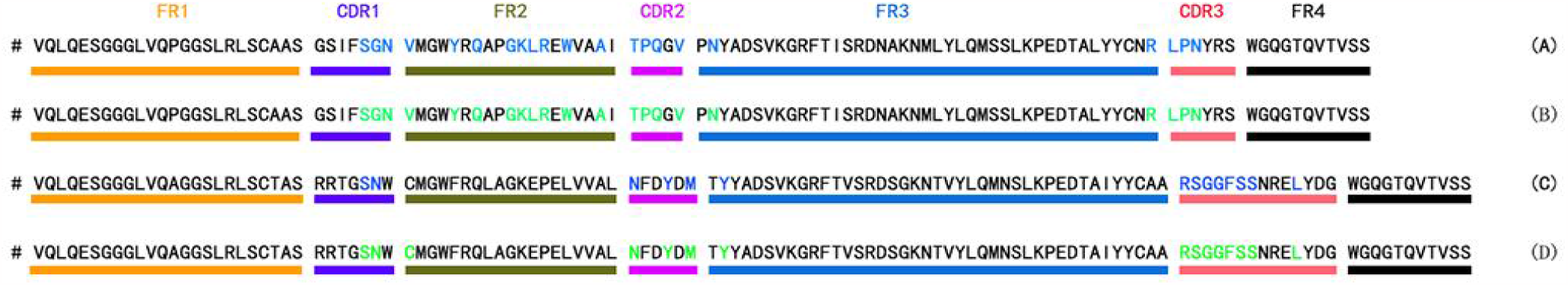
NanoBERTa-ASP accurately predicts the binding sites of nanobodies. (A) The computed binding sites from PDB:5f7y; (B) Prediction of PDB:5f7y binding sites by NanoBERTa-ASP; (C) The computed binding sites from PDB:2wzp; (D) Prediction of PDB:2wzp binding sites by NanoBERTa-ASP.

To verify the generalization ability of the model, we conducted 10-fold cross-validation on the model. We conducted tests using two different datasets: one consisting solely of nanobody sequences, and another consisting of the same number of heavy chain sequences added to the training set. To ensure that our evaluations were focused solely on the ability of the model to perform with respect to nanobodies, we used only nanobody sequences in our validation set. The AUC and precision obtained from the mixed dataset (AUC =0.952, precision=0.778) were higher than those from the pure nanobody dataset (AUC =0.947, precision=0.766). Through analysis of the results data, NanoBERTa-ASP has shown high stability in crossvalidation. [12]

We also compared our model with some currently available models for predicting binding sites, including ProtBERT, Paraperd, and Paragraph (Figure 4 and 5). As Paraperd and Paragraph only predict binding sites in the CDR region, we extracted the CDR part of the predicted results from the complete sequence predictions of ProtBERT and NanoBERTa-ASP for comparison.[13][14][15]

**Fig 4.**
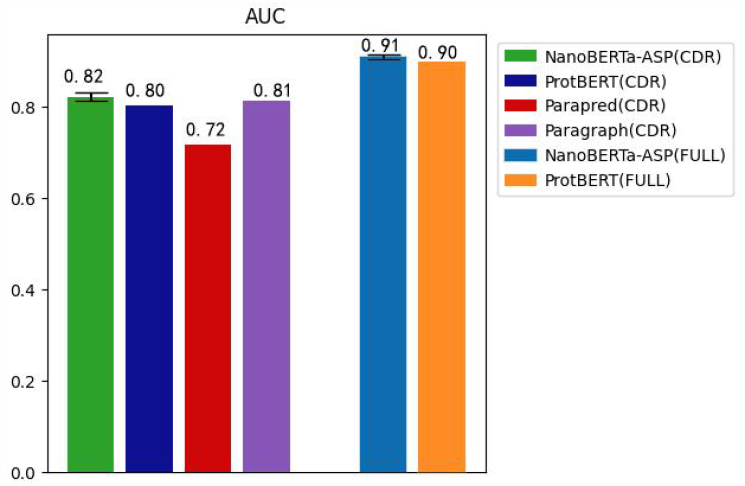
Comparison of NanoBERTa-ASP with Other Models Based on AUC Scores.

**Fig 5.**
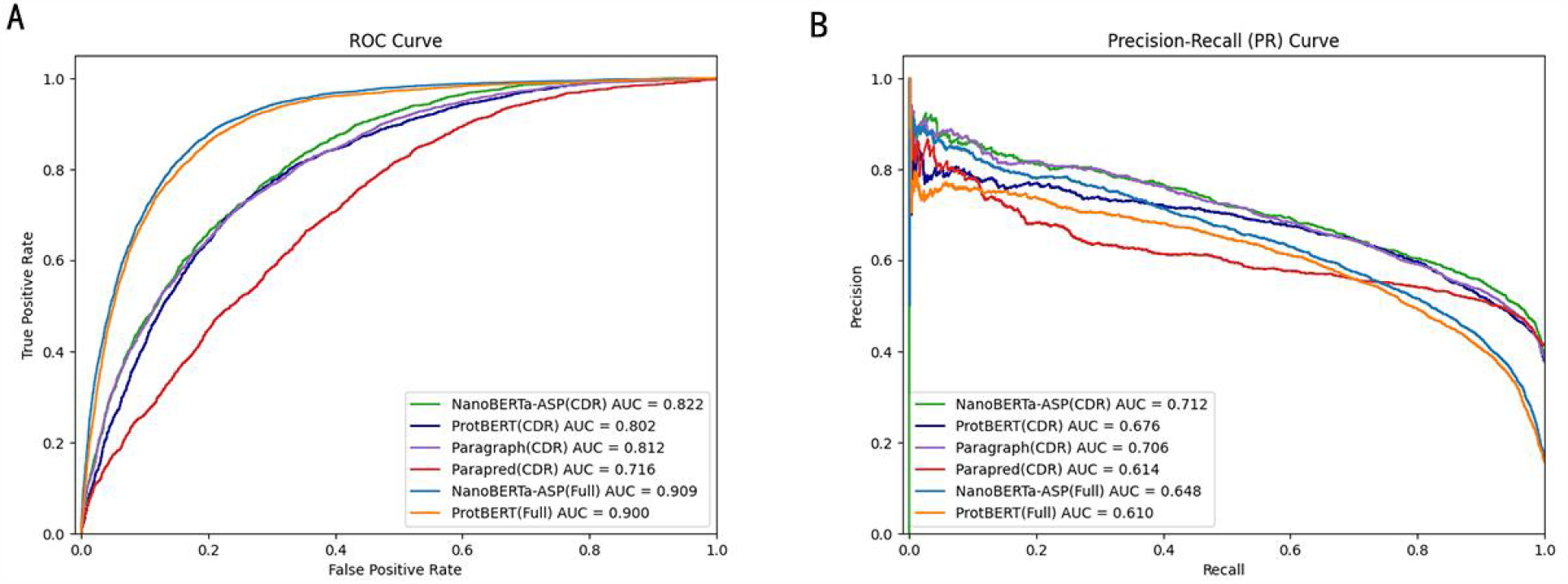
NanoBERTa-ASP outperformed other models in predicting binding sites Based on (A)ROC Curve and (B)Precision-Recall Curve.

NanoBERTa-ASP exhibits superior performance compared to publicly available tools in terms of the complete sequence of nanobodies and the CDR region. As the CDR region is a highly variable region that is harder to predict, and the main region where binding sites exist, the model is more focused on predicting positive samples, which may result in a lower AUC score for the CDR region than for the complete sequence, but a higher precision score in the CDR region (Figure 5). It demonstrated that NanoBERTa-ASP exhibits exceptional performance even with limited data, highlighting its significant potential for accurately predicting binding sites of nanobodies.

NanoBERTa-ASP, trained on a dataset of 30 million heavy chains, achieved comparable performance to ProtBERT, which was trained on a much larger dataset of 217 million proteins. This suggests that NanoBERTa-ASP is an effective model for nanobody sequence analysis, even with a smaller dataset.

## IV. CONCLUSION AND DISCUSSION

Regarding future improvements, for the dataset, we only performed simple screening. In future iterations of this research, clustering can be used to select heavy chains that are more similar to nanobody features for pre-training. Additionally, increasing the weight of the CDR region in the pre-training data can enable the model to maintain overall sequence attention while also directing greater attention to the highly variable regions, which may improve performance on downstream tasks. Furthermore, as the amount of data from cryo-electron microscopy- based nanobody experiments continues to increase, training with larger datasets and batch sizes could be explored in future studies.

For the nanobody binding site data used for fine-tuning, more accurate methods such as surface plasmon resonance and fluorescence spectroscopy can be used to obtain annotated information on binding sites. By improving the quality of the data, the model can learn the features of the binding sites more accurately.

